# Hidden endosymbionts: A male-killer concealed by another endosymbiont and a nuclear suppressor

**DOI:** 10.1101/2022.10.19.512817

**Authors:** Kelly M. Richardson, Perran A. Ross, Brandon S. Cooper, William R. Conner, Tom Schmidt, Ary A. Hoffmann

## Abstract

Maternally transmitted endosymbiotic bacteria that cause male killing (MK) have only been described from a few insects, but this may reflect challenges in their detection rather than a rarity of MK. Here we identify MK *Wolbachia* in populations of *Drosophila pseudotakahashii*, present at a low frequency (around 4%) in natural populations and previously undetected due to a different fixed *Wolbachia* strain in this species expressing a different reproductive manipulation, cytoplasmic incompatibility (CI). The MK phenotype was eliminated after tetracycline treatment that removed *Wolbachia*. Molecular analyses indicated the MK phenotype to be expressed when a second *Wolbachia* strain was present alongside the CI *Wolbachia*. A genomic analysis highlighted *Wolbachia* regions diverged between the strains involving 17 genes and also identified the *Wolbachia* as representing an outgroup to a clade of *Wolbachia* infecting *melanogaster*-group species, including *w*Ri-like and *w*Mel-like strains. Doubly infected males induced CI with uninfected females but not females singly infected with CI-causing *Wolbachia*. The MK phenotype manifested at the larval stage and was transmitted maternally at a high fidelity but with occasional loss of the MK *Wolbachia* strain. A rapidly spreading dominant nuclear suppressor genetic element affecting MK was identified through backcrossing and subsequent analysis with ddRAD SNPs of the *D. pseudotakahashii* genome. These findings highlight the complexity of nuclear and microbial components affecting MK endosymbiont detection and dynamics in populations, and the challenges of making connections between endosymbionts and the host phenotypes affected by them.

## Introduction

Male-killing (MK) phenotypes associated with endosymbionts were first investigated in ladybugs and butterflies [1]. While MK endosymbionts often occur at low frequencies in populations, they are expected to persist and spread when they provide a fitness advantage such as through the avoidance of sib mating [2, 3]. In *Drosophila*, several male-killers associated with *Wolbachia* and *Spiroplasma* endosymbionts have been described [4, 5]. However, their incidence in this genus is likely to be underestimated, partly because they can be uncommon in populations compared to *Wolbachia* that cause cytoplasmic incompatibility (CI) [6].

Male-killers in *Drosophila* typically result in embryo death; this includes male killing associated with both *Spiroplasma* [7] as well as *Wolbachia* [5, 6, 8] endosymbionts. Typically, such male-killers are detected by a reduction in hatch rate coupled with changes in sex ratio; this can be one reason for their underappreciation in natural populations given that male-killers are not maintained in stocks when males are required to produce offspring [9]. However, while reproductive effects of *Wolbachia* involving CI and MK are typically mediated through effects on embryonic development which results in a loss of egg hatch, they may also be affected by sex-specific mortality later in development. For instance, in mites and thrips, CI associated with *Wolbachia* has been reported as involving post-embryonic mortality [10, 11].

Although male-killers can result in all-female broods, there is variability in sex-ratio effects in some MK systems. In *Drosophila innubila, Wolbachia* density can vary among females which in turn correlates with female-biased offspring ratios, an effect that also has an epigenetic component and could contribute to stability of this infected system [4, 5]. Moreover, while some male-killer phenotypes can be stable across long time periods with little resistance to them evolving over thousands of years [12], MK phenotypes associated with endosymbionts can also be suppressed by nuclear genes.

A well-documented example of MK suppression is in the butterfly *Hypolimnas bolina*, where nuclear suppression resulted in a male-killer evolving into a CI phenotype [13, 14]. In this case, a high frequency of MK in a population which persisted for many years [15] was expected to produce strong selection for a nuclear suppressor because of the fitness advantage of rare males required for offspring production [16]. Rapid recovery of male production for male-killers associated with endosymbionts has also been documented in other systems including lacewings [17] and planthoppers [18]. The genetic basis of nuclear suppression is still unclear although in *Hypolimnas bolina* it involves a single chromosomal region [19] and in suppression generated following lab-based hybridization between two *Drosophila* species it is polygenic [20].

In *Drosophila pseudotakahashii*, Richardson et al. [21] described a CI *Wolbachia* infection present at a high incidence in natural populations and causing strong CI; however, the CI was weaker in older males from which the infection could be absent despite its high density in all females. Here, we describe a second *Wolbachia* strain in *D. pseudotakahashii* that is present at a low frequency and occurs alongside the CI strain where it causes MK. Unusually, male death occurred only after embryo development and was modifiable through a nuclear gene that segregated in some laboratory lines where it increased in frequency to the extent that sex ratio reverted. We use molecular approaches to characterize the MK strain which differs for some genomic regions to the coinhabiting CI strain but is identical in other regions. We also identify the genomic region associated with nuclear suppression through segregating crosses using the newly sequenced *D. takahashii* genome [22]. Our findings raise the issue of whether male-killers have been much more common in natural populations than previously assumed given that they are unlikely to be detected in the presence of common CI infections and may be affected by nuclear suppressors.

## Results

### A rare *Wolbachia* strain causes female-biased sex ratios

We established 188 *D. pseudotakahashii* isofemale lines from collections in Nowra, south-eastern Queensland and northern Queensland, Australia. Of these, 3.72% (*N* = 7) were found to have only female F1 offspring, although this differed slightly between collection sites (Table 1). No female-biased lines were found in the collections in northern Queensland or Moorland.

**Table 1.** Percentage of female-biased lines in the iso-female lines set up from field populations of *D. pseudotakahashii*

| **Collection area** | ***N*** | **% female-biased** |
| --- | --- | --- |
| **New South Wales** | **37** | **2.79** |
| Nowra | 34 | 2.94 |
| Moorland | 3 | 0.00 |
| **South-eastern Queensland** | **128** | **4.69** |
| Mount Tamborine | 74 | 4.05 |
| Cedar Creek | 10 | 0.00 |
| Mount Glorious | 44 | 6.82 |
| **Northern Queensland** | **23** | **0.00** |

*N* is the number of isofemale lines.

Sequences of individuals from female-biased and non-female-biased lines using the *Pgi* and *CO1* nuclear and mitochondrial markers showed almost no variation while sequences of the *Ddc* nuclear marker showed only a small amount of variation with no pattern separating female-biased and non-female-biased lines (sequences submitted to Genbank (accession numbers OP290972 - OP290995). These results, in addition to morphological examination of occasional males emerging from the female-biased lines supported the conclusion that the female-biased lines are indeed *D. pseudotakahashii*.

Nucleotide sequences from multiple female-biased lines were obtained for the five *Wolbachia* MLST loci and *wsp* [23]. Sequences presented a series of double peaks interspersed with sections without double peaks. These patterns were identical for the forward and reverse primers and present for all primer types and samples. Upon investigation, the ‘background’ sequence was the same as the *w*Pse CI [21] strain while the double peaks presented evidence for a second strain sharing many bases in common with *w*Pse. We designed strain-specific primers and screened a subset of the isofemale lines (*N* = 111) using standard PCR. Only the female-biased lines amplified with the MK primers, suggesting that the lines with female-only offspring were indeed the only lines with the double infection. Further genomic analyses (outlined below) confirmed the presence of two *Wolbachia* strains.

Treating copies of the female-biased lines with tetracycline resulted in emergence of male progeny and sex ratios that were closer to 50:50 compared to copies of the lines that were not treated with tetracycline (Table 2). RT-PCR with *wsp_validation* primers confirmed their uninfected status, and these lines became self-sustaining and no longer required the introduction of males from other lines, suggesting that the female-bias is indeed related to *Wolbachia* infection.

**Table 2.** Sex ratios of female-biased lines over the first 10 generations compared to non-female-biased and tetracycline-cured lines where *N* is the number of flies scored. *** denotes a deviation from a 50:50% male:female with chi-square tests (***: P < 0.001).

| Line | Phenotype | F1 | F4 | F10 |
| --- | --- | --- | --- | --- |
| | | | % Females ($N$) | % Females ($N$) |
| *B246<sup>MK</sup>* | female-biased | female-only | 99.19*** (143) | 54.41 (136) |
| *B256<sup>MK</sup>* | female-biased | female-only | 93.57*** (311) | 67.92*** (159) |
| *B280<sup>MK</sup>* | female-biased | female-only | 86.01*** (193) | 60.00 (95) |
| *B289<sup>MK</sup>* | female-biased | female-only | 87.17*** (226) | 56.30 (135) |
| *B302<sup>MK</sup>* | female-biased | female-only | 91.01*** (178) | 69.03*** (113) |
| *B305<sup>MK</sup>* | female-biased | female-only | 88.57*** (175) | 66.45*** (155) |
| *N101<sup>MK</sup>* | female-biased | female-only | 96.43*** (392) | 66.36*** (330) |
| *B116<sup>CI</sup>* | CI | males and females |  | 54.32 (81) |
| *N51<sup>CI</sup>* | CI | males and females |  | 48.48 (99) |
| *B302-* | tetracycline | N/A |  | 50.00 (88) |
| *B305-* | tetracycline | N/A |  | 46.88 (128) |
| *N101-* | tetracycline | N/A |  | 55.18 (299) |

### A double *Wolbachia* infection causes late MK

We assessed sex-ratio distortion by the *Wolbachia* double infection and its maternal transmission through crosses. We first crossed females from female-biased lines *N101*^*MK*^ and *B302*^*MK*^ with males from the non-female-biased *B116*^*CI*^ and *N51*^*CI*^ lines (*N* = 15). Three lines originating from *B302*^*MK*^ females produced both males and females (% female offspring of 16.67%, 20.00% and 42.86%), however the remainder of crosses had all female progeny. Female offspring from female-only lines (denoted by *‘MK1’*) and a line derived from *B302*^*MK*^ which produced male and female offspring (*‘MK2’*) were chosen for a second set of crosses. Note that we use italics to designate lines expressing MK and CI phenotypes associated with *Wolbachia*.

When *MK1* females were crossed with males from the *CI* lines (*B116*^*CI*^ and *N51*^*CI*^), uninfected line (*TPH35-*) or the mixed sex line (*MK2*), egg hatch proportions were similar to those involving females carrying CI-causing *Wolbachia* but the progeny were almost all female (Table 3). However, the mean percent egg-to-adult viability was half that of the control crosses. This suggests that male killing is occurring, but at a later time point than expected based on studies in other species (e.g. *D. pandora*, [6]).

**Table 3.**
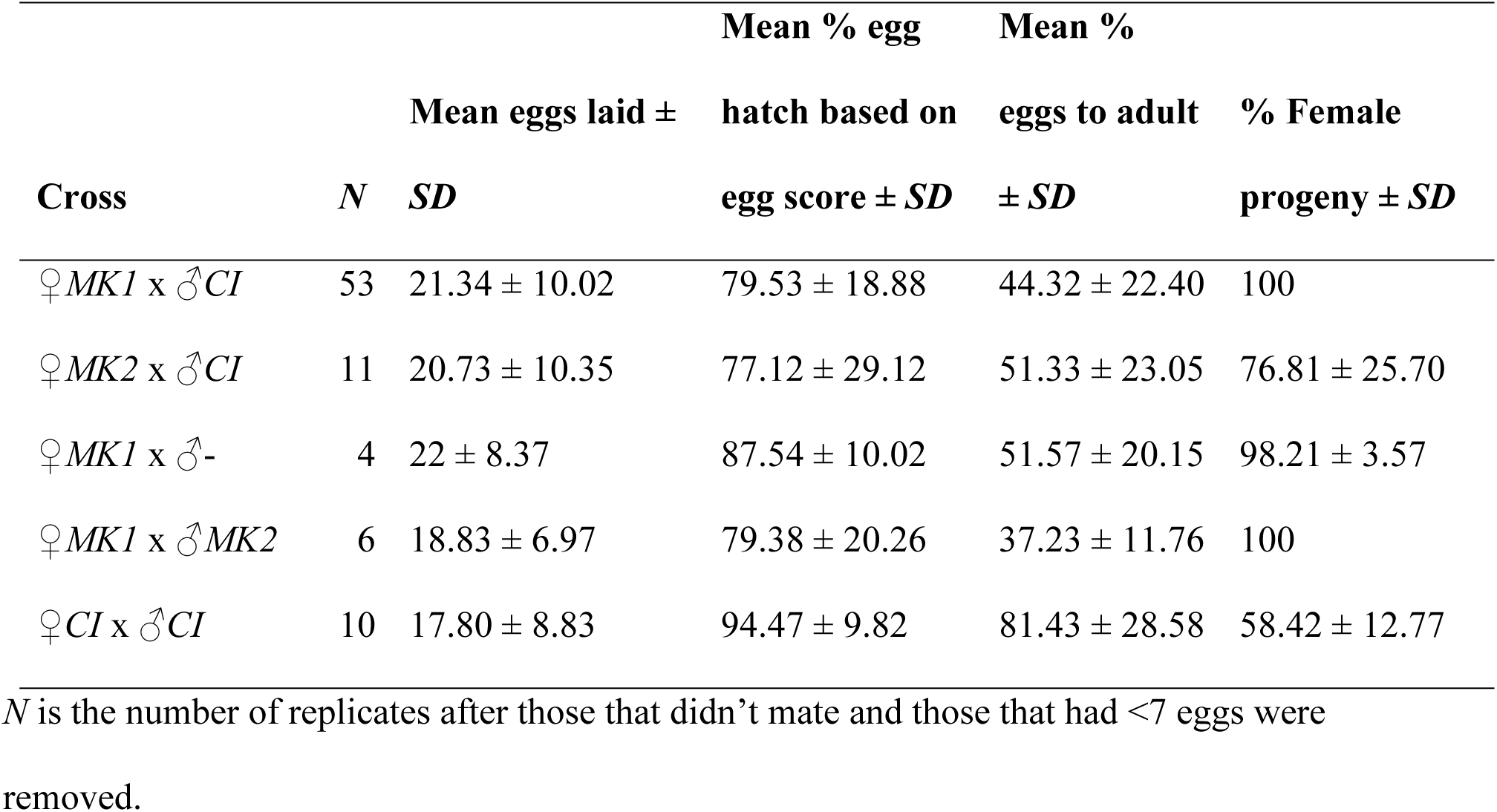
Crosses between individuals from *MK, CI* and uninfected (-) lines where *MK1* is from parents producing female-only offspring and *MK2* is from parents producing both male and female offspring.

When females from the mixed sex *MK2* line were crossed with CI males, males emerged from 5 of the 11 replicates (% female progeny of the 5 replicates ± *SD* = 53.63% ± 11.91) while the other 6 replicates had 100% female progeny, suggesting that the leakiness in MK in the female parent was not passed on to all her daughters. Mean % egg to adult viability (± *SD*) for the 5 crosses with males emerging was 68.31% ± 7.83 compared to 46.26 ± 13.00 for the replicates with only female progeny, indicative of MK rather than feminization.

Screening indicated that all but two females from the *N101*^*MK*^ and *B302*^*MK*^ lines used in the crosses had both the MK and CI *Wolbachia* strains (N = 73), suggesting incomplete maternal transmission, which is common in *Drosophila* [e. g. 24, 25]. Males from the *B116*^*CI*^ and *N51*^*CI*^ lines used in the crosses were 80% and 60% infected with the CI strain respectively (*N* = 45 and 25 respectively), consistent with findings elsewhere [21]. All *MK2* males had the CI strain (*N* = 9) while all but one had the MK strain.

### Rapid loss of MK, but long-term stability of MK *Wolbachia* during laboratory culture

Over time, males emerged in all the *MK* lines that were not tetracycline-treated (Table 2). By F10, the lines had only a slightly female-biased sex ratio. At F11, we tested two males and two females from a range of *MK* and non-female-biased lines and found that all but one female and all males from the female-biased lines carried the MK strain (Table S1). We tested a further 34 males from the *B289*^*MK*^, *B280*^*MK*^ and *B302*^*MK*^ lines (*N* = 7, 8 and 19 respectively) and all but two carried the MK strain, suggesting that despite the appearance of males, the MK infection was still present in the female-biased lines.

After 68 generations we again screened a subset of the lines for infection status. Despite the *MK* lines no longer having strong sex-ratio biases, all individuals tested from five *MK* lines were infected with the MK strain (females: *N* = 47, males: N = 47). Densities of the MK *Wolbachia* strain were similar between the sexes (GLM: F_1,84_ = 0.281, P = 0.597). Additionally, all females across the *CI* and *MK* lines were infected with the CI strain (*N* = 76). Consistent with previous research [21], presence of the CI strain was variable in males, with 90% of males from the *CI* lines (*N* = 30) and 89% of males from the *MK* lines (*N* = 47) carrying the CI strain, with *Wolbachia* density also being much lower in males than females (F_1,129_ = 145.531, P < 0.001). Sanger sequencing of the MLST genes from males singly infected with MK confirmed the presence of only the MK strain with alternate bases to the CI strain in locations where double peaks were present in double-infected individuals. Sequences have been submitted to Genbank (accession numbers OP290996 - OP291001).

### Genomic analyses point to a single CI and double MK *Wolbachia* infection

Table 4 presents assembly statistics for the *Smith+* and *N101*^*MK*^ draft *Wolbachia* assemblies, with the complete *w*Mel genome for comparison. *Smith+* assembled into a complete, circular genome. In contrast, we could not confidently separate the two *Wolbachia* infecting *N101*^*MK*^ or assemble a complete genome for either strain. The total number of genes meeting our criteria were similar for *Smith+* (*N* = 180), *N10*1^MK^ (*N* = 178), and for the *w*Mel reference genome (*N* = 180). BUSCO found 17 duplicated genes in *N101*^*MK*^, compared to zero in *Smith+* and in the *w*Mel genome.

**Table 4.** Assembly statistics for *Smith+* and *N101*^*MK*^, with the complete *w*Mel genome for

| Assembly Stats | *Smith+* | *N101<sup>MK</sup>* | *wMel* |
| --- | --- | --- | --- |
| Total Length | 1.31*10<sup>6</sup> | 1.57*10<sup>6</sup> | 1.45*10<sup>6</sup> |
| Number of scaffolds | 1 | 145 | 1 |
| BUSCO (Single\|Duplicated\|Fragment\|Missing) | 180\|0\|3\|38 | 161\|17\|3\|40 | 180\|0\|2\|39 |
| Longest scaffold | Complete | 107559 | Complete |
| N50 | Complete | 45215 | Complete |

Figures 1A and 1B show the normalized read depth of the *Smith+* and *N101*^*MK*^ lines, respectively, across the complete *Smith+* genome. As expected, *Smith+* has almost no deviations from 1 across the bulk of the genome, with the exception of four windows (a total of 3,000 bp) that have a normalized read depth over 3 (Figure 1B). This could plausibly represent sequencing bias or an assembly error in a small highly repeated area, since we do not expect CNVs when comparing *Smith+* to itself. In contrast, two regions of the *N101*^*MK*^ genome totaling approximately 150kb display approximately 0.7 normalized depth from positions around 600k to 700k and 900k to 950k. For 2->1 copy number changes, we would expect two paired regions, which we do not observe. While possible, for a potential 3->2 change (which would have depth close to 0.7) we would expect three affected regions. It seems more plausible that these genomic regions are present in the CI-causing *N101*^*MK*^ strain but not in the second *N101*^*MK*^ strain. There may be regions only present in the second strain, but our analysis would not identify those regions.

**Figure 1.**
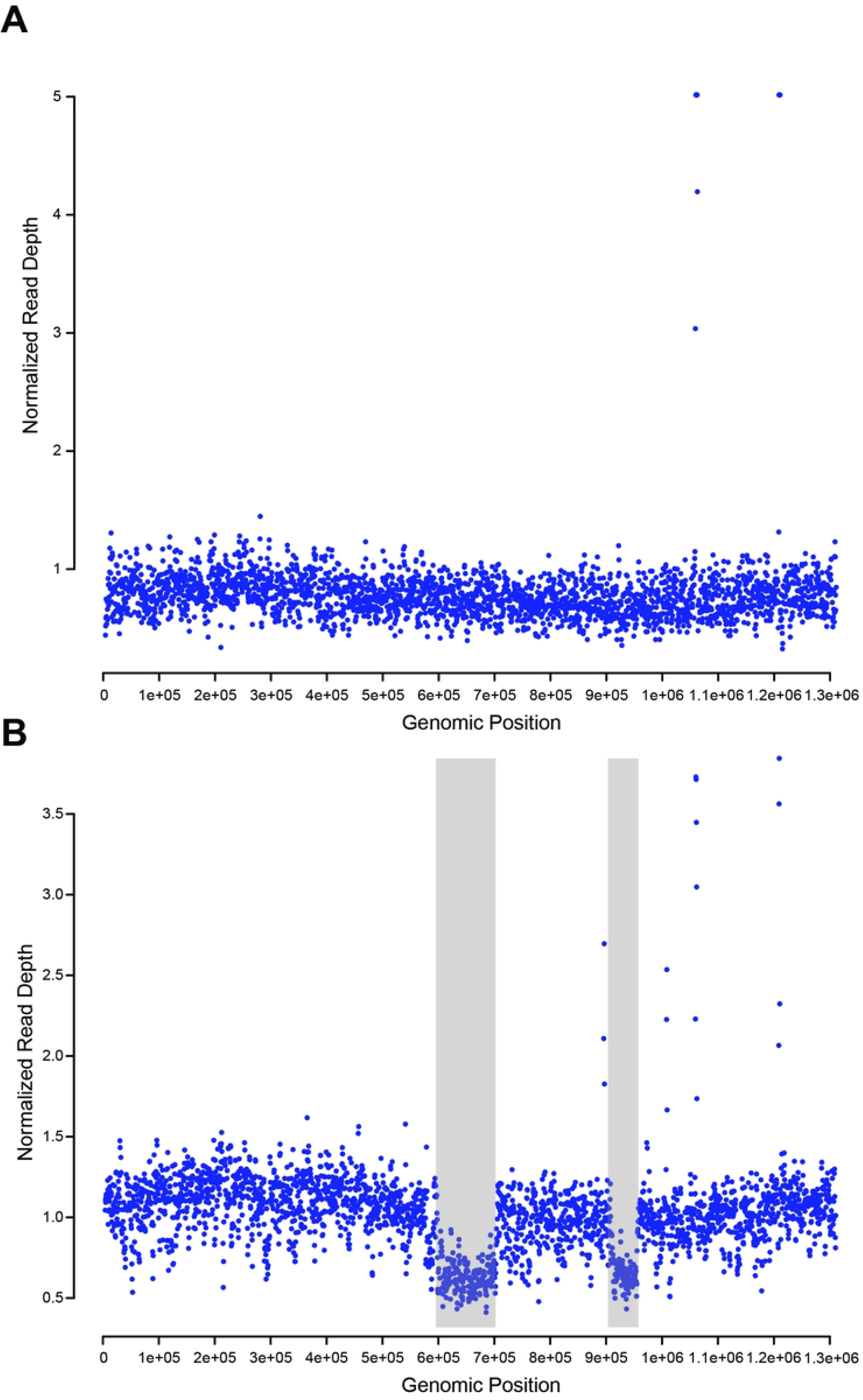
A) Normalized read depth of Illumina *Smith+* reads across the *Smith+* genome in 1000bp sliding windows. B) Normalized read depth of Illumina *N101*^*MK*^ reads across the *Smith+* genome in 1000bp sliding windows. Gray boxes denote regions displaying copy number variation. Depth was capped at 5 for readability.

Our analyses discovered CI-causing factors (*cifs*) in both the *Smith+* and *N101*^*MK*^ draft assemblies [26, 27]. *cifs* (*cifA/B*) are classified into five phylogenetic clades (Types I–V) [28, 29]. The *Smith+* genome contains one set of Type 1 *cif* loci and one set of the Type 2 loci. The *N101*^*MK*^ assembly contains two sets of Type 1 loci on a single contig and a single set of Type 2 loci on different contig.

Our analyses also discovered homologs of *w*Mel *wmk* in both the *Smith+* and *N101*^*MK*^ draft assemblies. There are two *wmk* copies in our *Smith+* assembly with 100% identity to each other and with 88.1% identity to *w*Mel *wmk* across bases 27-873; however, bases 1-26 and 874-912 across the region in *Smith+ wmk* have only 37% and 41% identity to *w*Mel *wmk*, respectively. The start and stop codons are intact as bases 1-3 and 910-912, respectively. In contrast, we found only one *wmk* copy in the *N101*^*MK*^ assembly, with 99.67% identity to *w*Mel *wmk* across all 912 bases. Notably, it remains unknown if transgenic MK produced by expressing *w*Mel *wmk* in *D. melanogaster* reflects anything about the genetic basis of MK in nature since *w*Mel does not cause MK in its natural *D. melanogaster* host, and has not been reported to cause MK in prior work introducing *w*Mel into other host species ([30]).

### Phylogenetic analysis shows links to the *w*Ri-like and *w*Mel-like groups

Our phylogram places the CI-causing *Wolbachia* that infects the *Smith+* genotype in Group A as an outgroup to a clade containing *w*Ha [31], *w*Ri-like (*w*Ri and *w*Ana [32]), and *w*Mel-like (*w*Au, *w*Mel, and *w*Yak [33]) *Wolbachia* (Figure 2). The four *Wolbachia* that infect *Nomada* bees (*w*NFe, *w*NPa, *w*NLeu, and *w*NFa [34]) are outgroup to the clade containing *w*Pse *Smith+*. These Group-A *Wolbachia* diverged from Group-B strains like *w*Pip_Pel in *Culex pipiens* [35] and *w*No in *D. simulans* [31] up to 46MYA [36]. We could not estimate the placement of the *Wolbachia* infecting the *N101*^*MK*^ genotype since we could not confidently separate the two strains.

**Figure 2.**
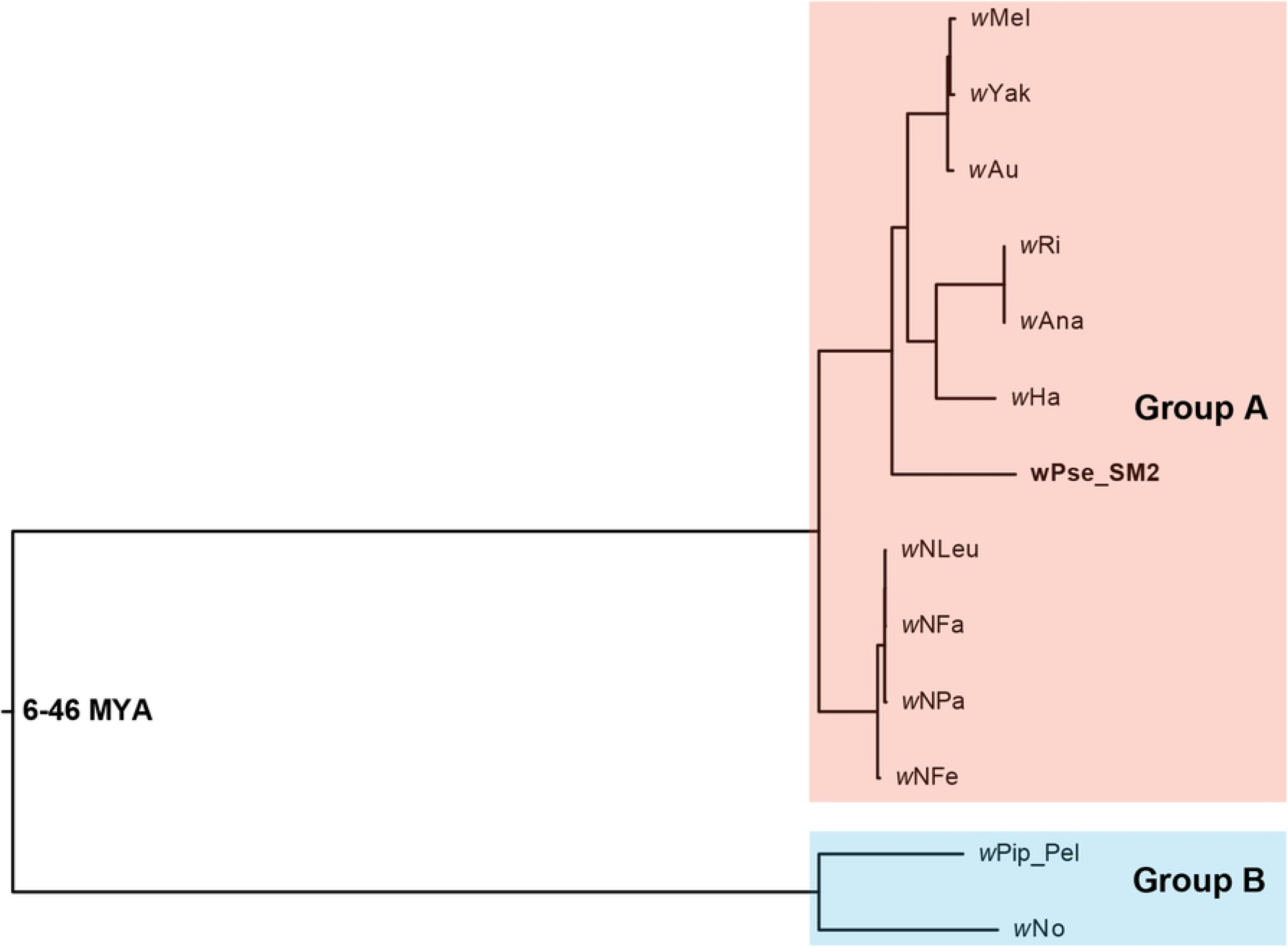
An estimated Bayesian phylogram for various Group-A (red) and Group-B (blue) *Wolbachia* strains. *w*Pse *Smith+* is a Group-A strain and outgroup to a larger clade containing *w*Ha, *w*Ri-like, and *w*Mel-like strains. The four *Wolbachia* infecting *Nomada* bees (*w*NFe, *w*NPa, *w*NLeu, and *w*NFa) are outgroup to the clade containing *w*Pse *Smith+*. These Group-A *Wolbachia* diverged from Group-B *Wolbachia* (*w*Pip_Pel and *w*No) up to 46MYA (divergence time superimposed from Meany et al. [36]). The phylogram was estimated with 168 genes and a total of 136,545 bp. Nodes with posterior probability <0.95 were collapsed into polytomies.

### Dominant MK suppression not linked to *Wolbachia* density

We introgressed two lines which maintained the MK *Wolbachia* strain but had no female bias (*N101*^*MKS*^ and *B302*^*MKS*^, with “*S*” in *MKS* denoting suppression of the MK phenotype) into the genetic background of a CI-only line treated with tetracycline (*B116-*) to test if the MK phenotype could be restored. Three out of 19 lines from the Nowra background reverted to MK (producing only females) after a single cross to *B116-* (Table 5), with the proportion of lines inducing MK increasing with subsequent backcrossing. Lines that produced only females continued to show a strong female bias in the following generations, with 93.75% (N = 32) of lines in backcross 3 and 92.31% (n = 39) in backcross 4 being female-only, with those producing males being highly female-biased (mean 90% female, N = 5). Crosses between females from the *MK* lines and males from the *MKS* lines produced only females, but their offspring included both sexes (Table 5). These results suggest dominant nuclear suppression of the MK phenotype.

**Table 5.** Segregation of MK suppression through crosses involving females with the MK *Wolbachia* strain that either expressed or did not express the MK phenotype.

| Female line | Males in backcross | Percent of lines with female-only offspring<br>(N female parents tested) |  |  |
| --- | --- | --- | --- | --- |
|  |  | F1 | Backcross 1 | Backcross 2 |
| *N101<sup>MKS</sup>* | *B116<sup>-</sup>* | 0 (20) | 15.79 (19) | 48.28 (29) |
| *B302<sup>MKS</sup>* | *B116<sup>-</sup>* | 0 (19) | 0 (20) | 39.29 (28) |
| *N101<sup>MKS</sup>* | *N101<sup>MKS</sup>* | 0 (20) | 0 (19) | 0 (16) |
| *B302<sup>MKS</sup>* | *B302<sup>MKS</sup>* | 0 (18) | 0 (20) | 0 (16) |
| *B116<sup>-</sup>* | *B116<sup>-</sup>* | 0 (20) | 0 (20) | 0 (17) |
| *N101<sup>MK</sup>* | *N101<sup>MKS</sup>* | 100 (9) | 4.76 (21) | 0 (17) |
| *B302<sup>MK</sup>* | *B302<sup>MKS</sup>* | 100 (9) | 0 (20) | 0 (17) |

To test whether MK suppression was associated with changes in *Wolbachia* density, we measured the density of the CI and MK strains in the *MK* (female-only offspring) and *MKS* (mixed-sex offspring) lines resulting from backcrossing as well as the original *N101*^*MKS*^ and *B302*^*MKS*^ lines (Figure S2). We found no difference in MK *Wolbachia* density between females from the *MK* and *MKS* lines (GLM: Nowra, F_1,28_ = 0.132, P = 0.719, Brisbane, F_1,28_ = 2.000, P = 0.168), indicating that suppression of the MK phenotype is not due to a decrease in *Wolbachia* density.

We performed crosses between *MK, MKS, CI* and uninfected lines to reassess the MK phenotype and test the ability of *MKS* males to induce cytoplasmic incompatibility. The offspring of *MK* females were strongly female-biased (Table 6), with only 2/22 replicates producing males. Despite similar egg-hatch proportions from crosses involving *MK* and *MKS* females, egg-to-adult viability of the offspring from *MK* females was half that of the other crosses. *MKS* males induced strong CI with uninfected females, with 6.17% of eggs hatching compared to ≥ 68% in the controls. Egg-hatch proportions in the *MKS* male x *CI* female cross were similar to the controls (Table 6), suggesting that the MK *Wolbachia* strain does not induce CI or has the same compatibility type as the CI strain.

**Table 6.** Crosses between individuals from *MK, MKS, CI* and uninfected (-) lines where *MK1* is from parents producing female-only offspring and *MK2* is from parents producing mixed-sex off spring.

| Cross | N | Mean % egg hatch | Mean % eggs to | % Female |
| --- | --- | --- | --- | --- |
| | | based on egg score ±<br>*SD* | adult ± *SD* | progeny ± *SD* |
| ♀*MK* x ♂*MKS* | 22 | 85.86 ± 7.33 | 32.22 ± 19.89 | 96.67 ± 11.55 |
| ♀*MKS* x ♂*MKS* | 27 | 73.54 ± 28.04 | 66.04 ± 33.34 | 44.94 ± 20.36 |
| ♀*MKS* x ♂*CI* | 27 | 73.70 ± 21.47 | 66.48 ± 33.19 | 53.92 ± 24.11 |
| ♀*CI* x ♂*MKS* | 25 | 82.71 ± 15.68 | 62.50 ± 26.35 | 44.44 ± 21.69 |
| ♀- x ♂*MKS* | 24 | 6.17 ± 8.14 | 62.84 ± 35.50 | 50.55 ± 35.44 |
| ♀*CI* x ♂*CI* | 20 | 68.00 ± 23.49 | 70.29 ± 35.75 | 46.01 ± 17.97 |
| ♀- x ♂- | 20 | 76.00 ± 24.10 | 60.72 ± 29.84 | 54.06 ± 19.52 |

*N* is the number of replicates after those that didn't mate and those that had <7 eggs were removed.

### Molecular analysis of segregating lines highlights a region with a selective sweep

We performed ddRADseq on lines derived from *N101*^*MK*^ and *B302*^*MK*^ which produced mixed-sex offspring (*MKS*) or female-only offspring (*MK*) to identify genomic regions associated with MK suppression (Figure 3a). We identified three contigs of >1 Mbp length where SNPs were structured in line with MK-suppression phenotype (Fig 3b). Of these, a specific region on contig NW_025323476.1 (positions: 3,321,074 – 4,677,392) showed strongly reduced variation in the *MK* lines but normal patterns of variation in the *MKS* lines, which matched expectations of a selective sweep on the *MK* lines. Genome-wide heterozygosity was lower in *MK* lines than in *MKS* lines (Table S2), though analysis of each contig in isolation showed that this difference was wholly due to heterozygosity differences in the three contigs from Figure 3B, where H_O_ was 23% smaller than other contigs in *MK* lines but 49% larger than average in *MKS* lines. Backcrossing was expected to produce negative genome-wide F_IS_, and this was more negative in *MKS* lines than *MK* lines. In the three contigs from Fig 3B, F_IS_ was 84% less negative than other contigs in *MK* lines but 60% more negative in *MKS* lines.

**Figure 3.**
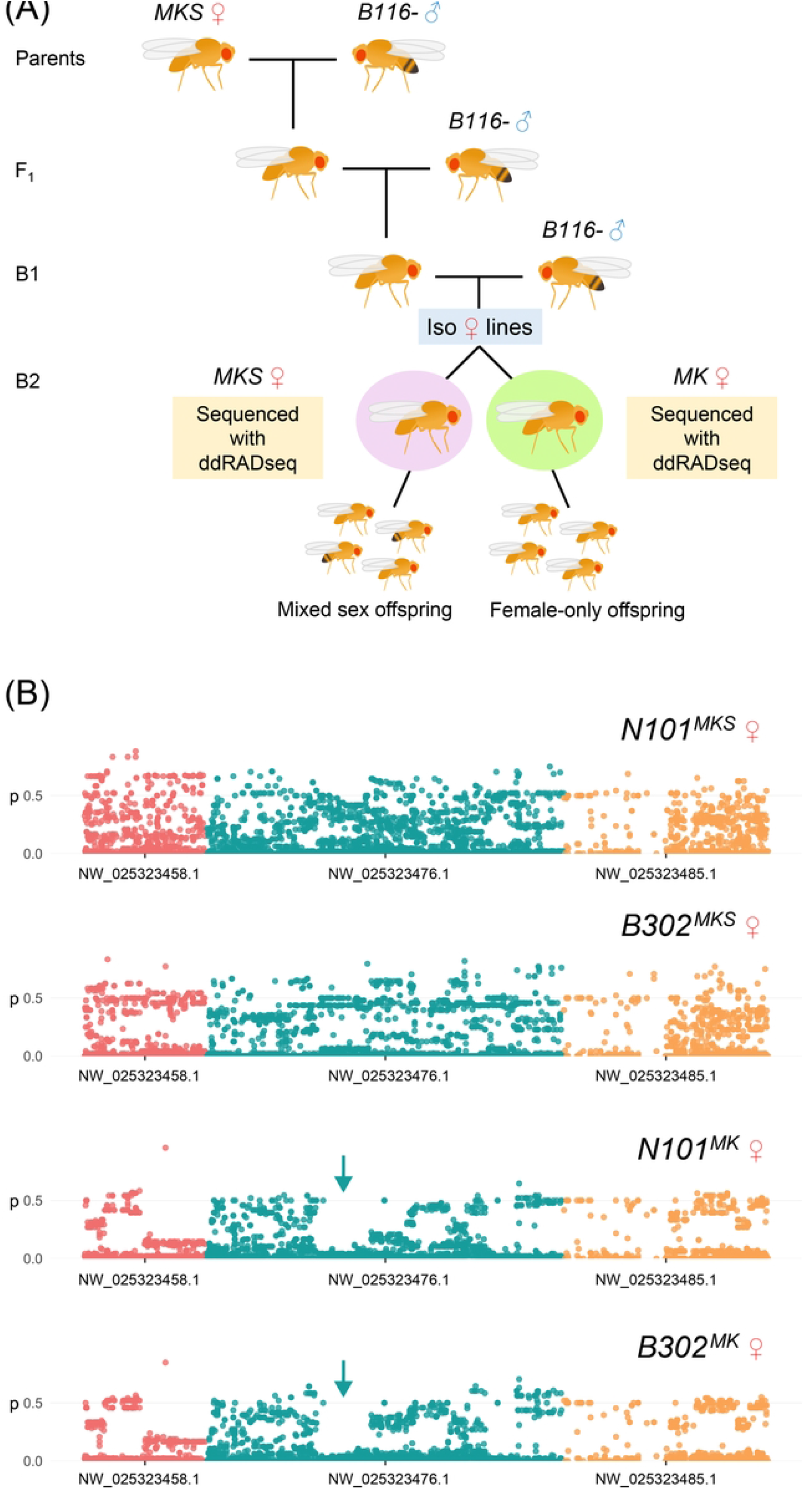
Molecular analysis of segregating lines. (A) is a cartoon detailing the experimental crosses. (B) shows frequency plots of non-reference alleles on the three contigs where structure followed MK suppression phenotype. A selective sweep pattern denoted by arrows is apparent on the NW_02532476.1 contig.

### MK suppression can spread rapidly in mixed populations

We set up mixed populations of *MK* (female-only offspring) and *MKS* (male and female offspring) females to track the spread of MK suppression across generations. Populations with only *MKS* females showed a relatively stable 1:1 sex ratio across generations, while the populations with only *MK* females remained stably and strongly female-biased (Figure 4). The mixed populations all tended towards a 50:50 sex ratio after a few generations, reflecting the strong spreading ability of the suppression genetic construct. Even the population that started with 90% *MK* reverted to a 50:50 sex ratio after only a few generations. To test for any fitness benefit that might be provided to female offspring by MK *Wolbachia*, we correlated sex ratio in experimental vials to the number of females produced in the following generation and found negative associations (G1, r = -0.304, P = 0.057, N = 40; G2, r = - 0.804, P < 0.001 N = 40). These results indicate that MK does not provide a detectable fitness benefit in terms of female production.

**Figure 4.**
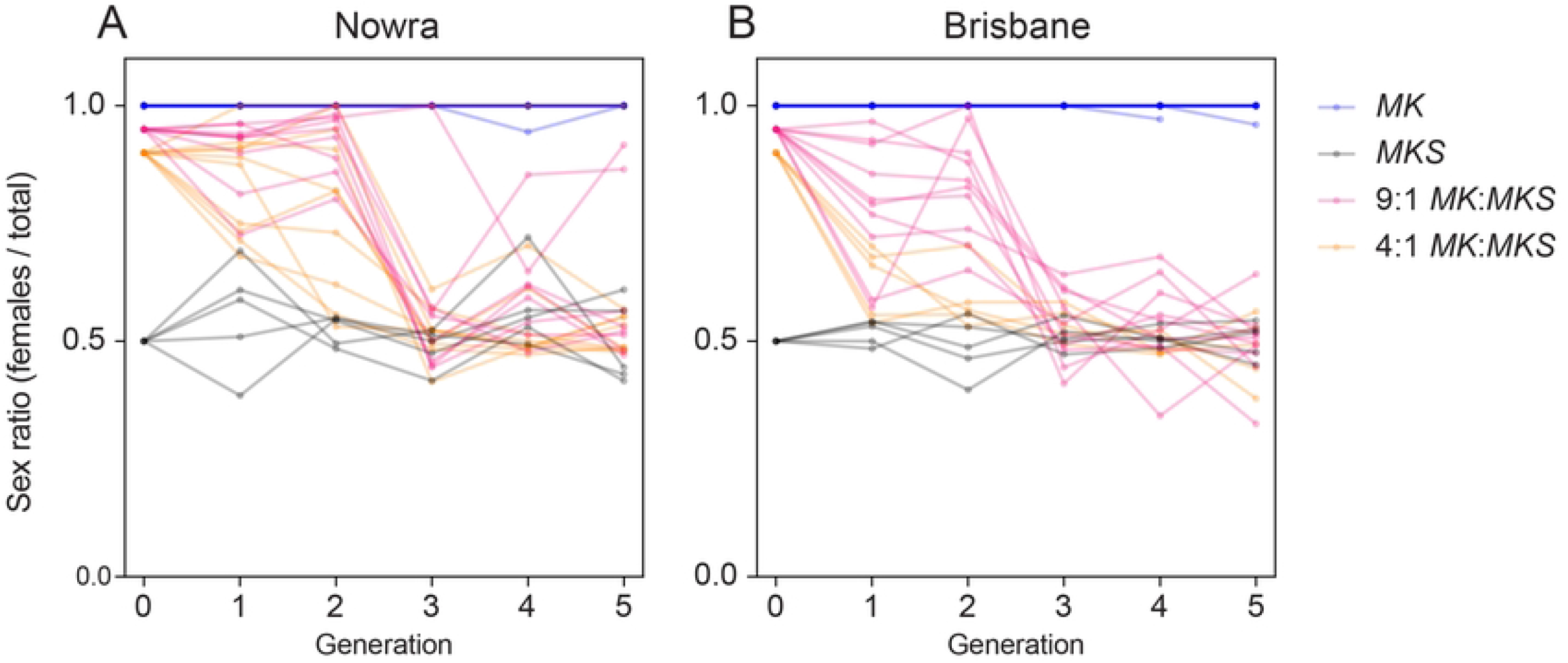
Changes in sex ratios across generations at different initial frequencies of *MK* and *MKS* females for the Nowra (A) and Brisbane (B) populations. Each line shows the sex ratio of a single replicate vial across generations Generation 0 denotes expected sex ratios based on starting ratios of *MK*:*MKS* females.

## Discussion

### A double Wolbachia infection associated with male killing

We report the presence of a double *Wolbachia* infection in *D. pseudotakahashii*, with the second strain inducing MK over and above the CI phenotype associated with the first strain. In previous studies on *Drosophila Wolbachia* infections, strains have been isolated that cause either CI or more infrequently MK, but in these cases the phenotypes are associated with strains that occur in separate hosts (e.g. [6]). Here we have evidence for two strains in the same individual. This was initially based on patterns of double peaks we observed in the original MLST/*wsp* analysis supported by the subsequent genomic analysis and the *gatB* MLST sequences that were used to design primers that could distinguish the strains.

In the genomic analysis, the draft *N101*^*MK*^ assembly we obtained was larger than the draft *Smith+* genome by approximately 260k bases, with BUSCO finding 17 genes duplicated in *N101*^*MK*^ compared to zero in *Smith+*. While a large novel insertion or duplication in *N101*^*MK*^ could potentially explain this, it seems more plausible that this pattern results from a double *Wolbachia* infection in the *N101*^*MK*^ genotype. If we assume that two closely related *Wolbachia* infect *N101*^*MK*^, as supported by preliminary analysis of MLST loci and by the normalized read depth plots across our draft genomes, similar regions sequenced from two unique genomic sources (i.e., two different *Wolbachia*) could have assembled into one scaffold. This would explain why the *N101*^*MK*^ assembly is only 260k bases longer at 1.57 million bp while a complete *Wolbachia* genome is typically on the order of 1 to 1.4 million bp.

This does not refute the insertion or duplication hypothesis, but a copy number decrease like 2->1 or 3->2 would require multiple identical regions to be affected on the normalized depth plots. For example, 2->1 would require two paired regions each at 0.5 depth and 3->2 would require three regions at 0.66 depth, which we do not observe. Rather, the 0.7 normalized average depth could be explained by titer differences between the two strains. If the CI-causing *Wolbachia* in *N101*^*MK*^ makes up 70% of the total *Wolbachia*, we would expect the regions only present in it to have depth of 0.7 times the rest of the genome. Future work aimed at separating these *Wolbachia* in doubly infected genotypes, or at identifying genotypes singly infected with the currently uncharacterized strain, will help confirm this.

### An uncommon infection that is nevertheless widespread

In *D. pseudotakahashii*, as in *D. pandora* and other species, the MK phenotype can be relatively uncommon and only detected when many lines from the field are screened. Nevertheless, we found the MK phenotype at multiple locations across the range of *D. pseudotakahashii*, a species found in cooler habitats along eastern Australia such as high elevation sites and rainforests [37]. These findings highlight the challenges involved in accurately characterizing *Wolbachia* phenotypes from molecular surveys where phenotypes are typically determined from only a few individuals and locations and where superinfections may not be recognized, particularly when *Wolbachia* titre can depend on temperature at both the high and low extremes [38, 39]. We suspect that superinfections and in particular combinations of CI and MK strains are more common than currently realized. There are currently relatively few cases where *Drosophila* species are superinfected although exceptions have been known for some time [e.g. 40]. Detection of an MK phenotype may also be less likely when MK phenotypes are only expressed at the larval stage. Crosses undertaken to characterize *Wolbachia* phenotypes may often be terminated at the egg stage when CI is normally expressed, including for CI expressed by *w*Ri group *Wolbachia* [32] which our genomic analysis shows to be closely aligned with the *w*Pse *Smith*+ strain.

The high level of the CI strain in *D. pseudotakahashii* is presumably maintained by strong CI and high maternal transmission. Although the CI *Wolbachia* density in males is low as evident in our study and also noted previously [21], CI was strong, consistent with the presence of CI loci in the *Smith+* genome. Our analyses revealed an additional copy of the Type 1 loci in *N101*^*MK*^ that we predict originates from a second *Wolbachia* in this putatively doubly infected genotype. Since both Type 1 loci reside on the same contig, it seems plausible this contig contains a misassembled chimera with elements from both strains.

Based on the lack of complete transmission of the double infection as detected in our experiments (unlike the CI strain), it is hard to see how the MK phenotype would be maintained in a population unless there is a fitness advantage to MK under some situations. This advantage may not need to be large, given that MK females are fully compatible with the CI inducing males. The MK infection remains stable in many laboratory lines after long term culture, even when the MK phenotype is lost. Moreover, there is no interaction between the density of the CI and MK strains in that the CI density is similar regardless of whether a line is also carrying an MK strain, suggesting that these strains are independent, which is further supported by the presence of a few males which have lost a detectable CI strain but where the MK strain was still found.

### A rare case of MK suppression

Few cases of MK suppression have been documented [41], and these have largely been uncovered through introgression experiments and transinfections. Here we observed the rapid spread of MK suppression when female-biased lines were brought into the laboratory, but find no evidence of MK suppression in natural populations. It is possible that both the expression of MK and suppression are environmentally-dependent, or that the advantages provided by MK are not selected for under laboratory conditions, such as when there is a strong level of sib competition which could favour the MK phenotype [42]. This situation contrasts with MK in another *Drosophila* (*D. innubila*) where no resistance to MK appears to have evolved over thousands of years [43].

In our study, MK was restored through introgression into a background with no suppression alleles, but suppression spread quickly when the suppression allele was present at low frequencies in mixed populations. This is not surprising because nuclear suppression alleles would be spread by both males and females – given that female *D. pseudotakahashii* with MK *Wolbachia* still need to mate, and offspring would then acquire suppression alleles that show dominance based on our backcrossing results. Although we have yet to identify the gene(s) involved, we have made progress in locating it to a chromosomal region where there are several candidates which now require finer scale mapping.

### Concluding remarks

Although the diverse phenotypic effects of *Wolbachia* and other endosymbionts have been recognized for some time [44], we have only recently started to make progress in understanding the population dynamics of multiple endosymbiont strains within the same individual and the interplay between the phenotypic effects associated with endosymbionts and the nuclear genome. We show here that this is a rich area for further analyses, particularly when coupled with recent advances in *Wolbachia* genomics and understanding the *Wolbachia* genes generating the phenotypic effects. Building this understanding is particularly important as endosymbionts start to be used in applied contexts where their evolutionary stability becomes critical.

## Methods

### *D. pseudotakahashii* field collections and laboratory lines

Female *D. pseudotakahashii* were obtained from two locations in New South Wales, three locations in south-eastern Queensland, and six locations in northern Queensland [21] (Table S3). All females were used to initiate isofemale lines. These collections yielded 7 female-biased (MK) lines from several south-east Queensland sites and a New South Wales site that were initially unidentifiable due to the complete absence of male progeny but suspected to be *D. pseudotakahashii*. By introducing males from identified lines of *D. pseudoakahashii*, we determined that the female-biased lines were *D. pseudotakahashii*. Species identification was based on occasional male offspring identified through sex combs on tarsomeres I and II of the male foreleg, and of the male terminalia [see 45]. Lines were screened for *Wolbachia* infection by PCR and RT-PCR (below).

To confirm that the female-biased lines were indeed *D. pseudotakahashii* and not a cryptic species, we used the *Drosophila* nuclear markers *Ddc* and *Pgi* [46] and the mitochondrial marker *CO1* [47] to screen a single individual from three confirmed *D. pseudotakahashii* lines that induce CI (*B116*^*CI*^, *N51*^*CI*^ and *Smith+*) and five female-biased lines (*N101*^*MK*^, *B305*^*MK*^, *B302*^*MK*^, *B256*^*MK*^ and *B289*^*MK*^). DNA extractions using a 5% Chelex (Bio-Rad Laboratories, Gladesville, NSW, Australia; Cat. No. 142-1253; w/v in distilled water), and PCR conditions were performed as outlined in Richardson et al [21]. PCR products were Sanger sequenced by Macrogen (Korea) and chromatograms were checked and edited manually in Finch TV v1.4.0 (Geospiza, Seattle, WA) and MEGA version 6 [48]. Sequences have been deposited in Genbank (accession numbers OP290972 – OP290995).

Lines of *D. pseudotakahashii* used in crosses and experiments (Table S3) were female-biased lines *N101*^*MK*^ and *B302*^*MK*^ and non-female biased lines *N51*^*CI*^ and *B116*^CI^. Females from all lines were infected with *Wolbachia* based on PCR characterization (see below). Because there were no naturally uninfected lines [see 21], uninfected lines were generated for crosses from the isofemale line *TPH35*^*CI*^ *(Town3*^*+*^) by treatment with 0.03% tetracycline (Sigma, Castle Hill, NSW, Australia) in cornmeal media for one to two generations (as outlined in Hoffmann et al [49]). The derived *Wolbachia*-negative lines are designated with a “^−^” symbol as *TPH35*^*-*^. Three of the female-biased lines (*N101*^*MK*^, *B302*^*MK*^ and *B305*^*MK*^) and one non-female biased line (*B116*^*CI*^) were also treated with tetracycline as outlined above and their treated counterparts denoted by *N101*^*-*^, *B302*^*-*^, *B305*^*-*^ and *B116*^*-*^. The sex ratio of progeny was scored after curing and the removal of *Wolbachia* infection was also verified via RT-PCR (see below). Lines were maintained in the laboratory on cornmeal media at 19°C with a 12:12 L:D cycle and treated lines were allowed to recover in the absence of tetracycline for at least 2 generations before being used in experiments.

Lines (including those where the infection had been removed with tetracycline treatment) were monitored across time and their sex ratio was determined at F4 and F10 since setting up lines from the field. We collected at least 80 (range 80-390) per line to score sex ratio and compared results to an expectation of 50:50% male:female with chi-square tests.

### Crossing patterns and *Wolbachia* maternal transmission

We characterized the female-biased *D. pseudotakahashii* lines *N101*^*MK*^ and *B302*^*MK*^ by conducting a series of experiments investigating maternal transmission, CI and sex ratio distortion. Experiments were conducted at 19°C with a 12:12 L:D cycle.

We assessed maternal transmission of *Wolbachia* in *D. pseudotakahashii* female-biased lines in a two-part experiment using F4 and F5 individuals. In the first part, we crossed females from female-biased lines *N101*^*MK*^ and *B302*^*MK*^ with males from the *B116*^*CI*^ and *N51*^*CI*^ lines (*N* = 15). Of these, four lines were selected for part 2 of the experiment (including a line that produced both males and females denoted by *MKS* in which part 2 females were crossed with males from the CI lines *B116*^*CI*^ and *N51*^*CI*^ (*N* = 84) and males from the *MKS* line (*N* = 10) in addition to control crosses between the CI lines (*N* = 12). For comparison, we also included crosses between the *N101*^*MK*^ and uninfected *TPH35*^*-*^ males (*N* = 4), and uninfected females and *MKS* males (*N* = 4), however replicate numbers were not high enough to investigate trends.

Crosses were set up when individuals were 4-7 days old, mating was observed after which males were removed and stored in ethanol. Females were provided with spoons containing cornmeal media and a brush of yeast paste to encourage egg-laying. Spoons were scored for egg number and replaced every 24 hours for up to four days. Twenty-four hours after collection, eggs were scored for hatched and unhatched eggs. Given the comparatively low egg laying potential for this species [21], we used spoons where 7 or more eggs had been laid. Replicates that didn’t mate and had fewer than 7 eggs were removed from analysis. Progeny took approximately 18 days to develop, emerging adults were stored in 100% ethanol and sexed.

### Long-term stability of the MK *Wolbachia* infection

Sex ratios of the *N101*^*MK*^ and *B302*^*MK*^ lines were scored approximately 68 generations after the lines were initiated. We also reassessed infection status and determined the infection densities for 10 males and 10 females from a subset of the lines (originally female-biased: *N101*^*MK*^, *B305*^*MK*^, *B302*^*MK*^, *B289*^*MK*^, *B256*^*MK*^ not female-biased: *Smith+, N51*^*CI*^, *B116*^*CI*^) using the Roche LightCycler^®^ 480 system and the strain-specific qPCR primers outlined below.

### *Wolbachia* strain detection and strain typing

A preliminary screen for *Wolbachia* infection was conducted for all field isofemale lines. DNA extractions were performed using the 5% Chelex based method outlined above. Samples were screened for *Wolbachia* via RT-PCR using the *wsp_validation* primers [50, 51]. This assay determines *Wolbachia* infection from the melting temperature (T_m_) of the *wsp* validation PCR amplicons. High-resolution melt analysis on the Roche LightCycler^®^ 480 system produced a T_m_ range from approximately 81.1 – 81.9 °C for *D. pseudotakahashii*, including the female-biased lines.

To investigate the *Wolbachia* infection of the female-biased lines in more detail, we initially used the forward and reverse *coxA, hcpA, ftsZ, fpbA* and *gatB* MLST primers [23] and *wsp_validation* primers [50, 51] to screen a single individual from *N101*^*MK*^, *B302*^*MK*^ and *B289*^*MK*^. Conditions were as outlined in Richardson et al [6], PCR products were sent to Macrogen (Korea) for purification and Sanger sequencing chromatograms were examined and processed as outlined above using Finch TV v1.4.0 (Geospiza, Seattle, WA) and MEGA version 6 [48]. Sequences revealed the presence of double peaks for many of the MLST markers - notably absent in chromatograms of the *D. pseudotakahashii Wolbachia* strain *w*Pse generated by Richardson et al [21]. Double peaks can be indicative of double infection [23] and although we initially used MLST as an established system which would allow links to be made to previous work, we conducted further sequencing analysis of the potential double infections using whole genome sequencing, which is a preferable approach [32, 52].

For general screening of field and experimental samples we were unable to use established genotyping assays that distinguish CI and MK strains on the basis of melt analysis using *wsp_validation* primers (e.g. Richardson et al [6]) because the Tm of the two strains in *D. pseudotakahashii* overlapped (81.1 – 81.9 °C). Instead, we initially designed reverse primers to distinguish the two strains via standard PCR, based on alignments from the *gatB* MLST sequences we generated for the female-biased and non-female-biased lines (gatB_pt_MK_R: 5’-GTATCTATAATCGCTTGCATCCTC-3’ and gatB_pt_CI_R: 5’-GGCAACAAGTCAGGCTCA3’) and used these in conjunction with the existing *gatB* forward primers to produce bands of 373bp and 388bp that amplified the female-biased and non-female-biased strains respectively. The PCR conditions for both primers were the same (3 mins at 94°C; 37 cycles of 30 seconds at 94°C, 45 seconds at 71°C, 90 secs @ 72°C and a final extension for 10 mins at 72 °C followed by holding at 4°C). A subset of the isofemale lines initiated from field collections including 98 lines from Brisbane, 11 from far northern Queensland and 2 from Nowra were screened with these primers. To determine relative densities of the strains, we later used genomic sequences (outlined below) to design strain-specific primers specifically for use on the Roche LightCycler^®^ 480 system (MK strain: MK_F1; CCTTGTATTGAACTTCATCTTTGTTAC and MK_R1; GAACTTGTTTTACTTTATCACTTATCAC and CI strain: CI_Fc; TTCGATAAATAGACTTTTAAACTCTGTA and CI_Rc; TTTTAGACAATCTTGATAATCTTGC). Primer-specificity was confirmed using several samples that were known to be singly infected with CI or MK and ensuring there was no cross amplification with non-strain-specific primers. The qPCR conditions for the CIc and MK3 primers were the same with an annealing temperature of 63°C. Using this system, we generated crossing point (Cp) values for MK and CI strain-specific markers (outlined below) in addition to the *Drosophila* universal primers targeting the ribosomal protein gene *L40* (RpL40) [21]. Densities were calculated based on methods outlined in [21] using the mean Cp value generated by two replicate runs and a standard deviation threshold of 2.5.

### *Wolbachia* whole genome sequencing and assembly

We next sequenced whole genomes for the *N101*^*MK*^ and *Smith*+ genotypes using both short- and long-read technology. To produce short reads, total DNA was extracted from males (*N* = 5) and females (*N* = 5) of each genotype using the DNeasy Blood & Tissue kit following manufacturer protocol (Qiagen). Illumina libraries were prepared using the Nextera DNA Flex Library Preparation Kit (Illumina). Final fragment sizes and concentrations were confirmed using a TapeStation 2200 system (Agilent), and the samples were indexed using IDT for Illumina Nextera DNA UD indexes. Pooled libraries were shipped to Novogene (Sacramento, CA, USA) for sequencing on a partial Illumina NovaSeq lane, generating paired-end 150 bp reads. DNA was extracted from pooled individuals (*n* = 10) using a DNeasy® Blood & Tissue kit (Qiagen, Hilden, Germany). A continuous long-read library was prepared and sequenced using PacBio Sequel II technology by Berry Genomics (Berry Genomics Co. Ltd, Beijing, China). PacBio sequencing yielded 8.91E+05 reads from *N101*^*MK*^ and 8.22E+05 reads from *Smith*+, with a fasta size of 1.53E+10 and 1.57E+10 bp and approximate coverage of 50.96 and 133.65 The PacBio reads for the *N101*^*MK*^ and *Smith+* genotypes were assembled, and the *Wolbachia* contigs were extracted. Our assembly resulted in a final set of 314 scaffolds from *N101*^*MK*^ with total length of 1.58E+06 bp and N50 of 19011 bp, and 38 scaffolds from *Smith*+ with total length of 1.26E+06 bp and N50 of 62575 bp.We polished the PacBio assemblies using the Illumina libraries and pilon v 1.23 [53] set to default parameters.

To evaluate the quality of the draft *Wolbachia* assemblies, we used BUSCO 3.1.0 [54] to search for homologs of the single-copy genes in the proteobacteria database. As a control, we performed the same search using the reference *w*Mel genome [55].

### Genomic analyses

To assess the genomic overlap and potential copy number variants (CNVs) in *N101*^*MK*^ compared to *Smith*+, we aligned the Illumina reads for N101 and *Smith+* to the draft *Smith+ Wolbachia* genome using bwa 0.7.17 (Li and Durbin 2009 [56]). (The draft *Smith+ Wolbachia* genome was a single scaffold.) Normalized read depth for each alignment was calculated over sliding 1000 bp windows (1-1000, 500-1500, etc.) by dividing the average depth in the window by the average depth over the entire genome. The normalized read depth was plotted and visually inspected for regions with normalized depth different from 1. We capped the normalized depth of each window to 5 for readability.

Recent research has identified CI-causing factors (*cifs*) associated with WO prophage in *Wolbachia* genomes [26, 27, 57], including individual SNPs that influence CI strength [58]. We searched for *cifs* in *w*Pseu by BLASTing *cifs* from *w*Mel (Type 1 loci), *w*Ri (Type 2 loci), *w*No (Type 3 loci), *w*Pip (Type 4 loci), and *w*Stri (Type 5 loci) against our draft *N101*^*MK*^ and *Smith*+ assemblies.

While the genetic basis of MK remains unknown, the *wmk* gene associated with the WO prophage region of non-MK *w*Mel kills male embryos when transgenically expressed in *D. melanogaster* [59]. We specifically searched for *wmk* by BLASTing *w*Mel *wmk* (WD0626) against our draft *N101*^*MK*^ and *Smith*+ assemblies.

### Phylogenetics

To extract genes for our phylogenetic analyses and identify homologs to known bacterial genes, we annotated our draft *Smith+* genome and the public genomes of *w*Ana from *D. ananassae* [60]; *w*Au, *w*Ha, *w*Ri and *w*No from *D. simulans* [31, 61, 62]; *w*Mel from *D. melanogaster* [55]; *w*NFa, *w*NFe, *w*NLeu and *w*NPa from *Nomada* bees [34]; *w*Pip_Pel from *Culex pipiens* [35]; and *w*Yak from *D. yakuba* [33] with Prokka v.1.11 [63]. To avoid pseudogenes and paralogs, we used only genes that were present in a single copy and with identical lengths in all of the analyzed sequences. Genes were identified as single copy if Prokka uniquely matched them to a bacterial reference gene. By excluding homologs that were not of equal length in all of our draft *Wolbachia* genomes, we removed all loci with indels across any of the included sequences. In total, 168 genes with a combined length of 136,545 bp met these criteria. We did not include *Wolbachia* infecting the *N101*^*MK*^ genotype in our phylogenetic analyses since we could not confidently separate them.

With these 168 genes, we estimated a phylogram using RevBayes v. 1.1.1, following the procedures of Turelli et al. [32]. Briefly, we used a GTR + gamma model with four rate categories, partitioning by codon position. Each partition had an independent rate multiplier with prior Γ(1,1) [i.e., Exp(1)], as well as stationary frequencies and exchangeability rates drawn from flat, symmetrical Dirichlet distributions [i.e., Dirichlet(1,1,1…)]. The model used a uniform prior over all possible topologies. Branch lengths were drawn from a flat, symmetrical Dirichlet distribution, and thus summed to 1. Since the expected number of substitutions along a branch equals the branch length times the rate multiplier, the expected number of substitutions across the entire tree for a partition is equal to the partition’s rate multiplier. Four independent runs were performed and all converged to the same topology. Nodes with posterior probability <0.95 were collapsed into polytomies. For additional details on the priors and their justifications, consult Turelli et al. [32].

### Segregation of MK suppression

#### Generation of lines

We performed crosses to determine the basis of MK suppression and generate lines expressing MK or a 50:50 M/F sex ratio for molecular analyses. Unmated females from the *N101*^*MKS*^ and *B302*^*MKS*^ lines were mated individually to *B116*^−^ males, with 20 replicates per cross. The *N101*^*MKS*^ and *B302*^*MKS*^ lines (with “S” denoting suppression of MK) carried both the CI and MK *Wolbachia* strains but no longer showed any female bias, while the *B116*^−^ line was cured of its CI *Wolbachia* strain and was not previously infected with the MK strain. Offspring from each replicate vial were scored for sex ratio, then unmated female offspring were pooled across replicates and mated to *B116*^−^ males individually, with this process repeated for two backcrosses. Three replicates of the cross between *N101*^*MKS*^ and *B116*^−^ lines produced female-only offspring; we set up an additional 10 replicates of each line and continued to cross them to *B116*-males to test whether the female-only phenotype was maintained in the following generations. After backcross 2, we set up a third backcross, with separate sets of crosses with female-only lines and mixed sex lines, with 20 replicates each. Additionally, 60 replicate females each of the *N101* and *B302* lines from backcross 2 were isolated following mating within the line, then stored for molecular analysis after producing offspring (Figure 3A). In each generation 20 replicates each of the *N101*^*MKS*^, *B302*^*MKS*^ and *B116*^−^ lines (self-crossed) were set up as controls, with an expected 1:1 sex ratio each generation. In all crosses, flies allowed to mate for 3 days then transferred to new food to produce offspring for 4 days.

In a second experiment, we reverted female-only lines to produce both male and female offspring. Individual females from the female-only lines derived from *N101*^*MK*^ and *B302*^*MK*^ were crossed to males from the *N101*^*MKS*^ and *B302*^*MKS*^ lines respectively, with 10 replicates each. Female offspring were then individually mated to *N101*^*MKS*^ and *B302*^*MKS*^ males in the F1 and B1 generations, with 20 replicates per cross.

To test whether suppression of MK was associated with changes in *Wolbachia* density, we determined the infection densities of the CI and MK *Wolbachia* strains from backcrossed *N101*^*MK*^ and *B302*^*MK*^ lines that produced only females or both male and female offspring, as well as the original *N101*^*MKS*^ and *B302*^*MKS*^ lines. Fifteen females and 15 males (when present) from each group were screened using the Roche LightCycler^®^ 480 system and the strain-specific qPCR primers outlined below.

#### Reassessment of crossing patterns in segregated lines

We performed an additional set of crosses to test the ability of *B302*^*MKS*^ males to induce CI with *B116*^−^ and *B116*^*CI*^ females. We also tested whether the *B302*^*MK*^ line, which was reverted to MK as described above, induced late-acting MK. Crosses were performed as described above (see “Crossing patterns and *Wolbachia* maternal transmission”), with thirty replicates established per cross. We included crosses within each line for *B302*^*MKS*^, *B116*^*CI*^ and *B116*^−^ as controls, which were expected to show high egg hatch and egg to adult viability with a 50:50 sex ratio.

#### Molecular analysis of segregating lines

To investigate potential genes driving suppression of the MK phenotype we utilised the double-digest restriction-site associated DNA sequencing (ddRADseq) protocol developed by Rašić et al [64] to construct ddRAD libraries and genotype females for genome-wide SNPs. Females were those resulting from backcross 2 which were isolated and stored for ddRADseq following oviposition (see above). We separated females into two groups for each of the *N101* and *B302* lines; those that produced female-only offspring (*MK*), and those that produced both male and female offspring (<70% female, *MKS*). Lines producing >70% females or fewer than 15 total offspring were excluded due to phenotype ambiguity (since MK may be leaky). DNA from twenty-four females from each of the four groups was extracted using Roche High Pure™ PCR Template Preparation Kits (Roche Molecular Systems, Inc., Pleasanton, CA, USA). A total of 96 *D. pseudotakahashii* females from across the four groups were grouped into four libraries and sequenced by NovogeneAIT Genomics (Singapore).

Sequence data were processed with Stacks v2.54 [65]. We used the *process_radtags* program to demultiplex sequence reads, discarding reads with average Phred score below 20. We used bowtie v2.0[66] to align reads to the *D. takahashii* genome assembly GCA_018152695.1 [67] using --very-sensitive alignment. Genotypes were called with the *ref_map* program, with effective per-sample coverage of 24.5 ± 7.1X. We used the *Populations* program to calculate genome-wide heterozygosity and F_IS_ for each group, omitting sites with any missing genotypes (-R 1) and retaining both monomorphic and polymorphic sites [68]. We ran *Populations* again to output a set of SNPs that were called in ≥ 95% individuals and when ≥ 3 copies of the minor allele were present [69]. We visually compared allele frequency plots of the remaining 26810 SNPs for regions where genetic structure covaried with MK suppression across the *N101* and *B302* lines.

#### Spread of MK suppression in mixed populations

To investigate the spread of MK suppression, we set up mixed populations of *N101*^*MK*^ or *B302*^*MK*^ females (which produced female-only offspring) and *N101*^*MKS*^ or *B302*^*MKS*^ females (which produced both male and female offspring) and tracked sex ratios across generations. *MK* and *MKS* females were mated to *B116*^*-*^ and *N101*^*MKS*^*/B302*^*MKS*^ males respectively, then added to vials at ratios of 9:1 and 4:1 *MK*:*MKS*. We set up replicate vials of *N101*^*MKS*^, *B302*^*MKS*^ and *B116*^−^ as controls which were expected to maintain a 50:50 sex ratio across generations. To test whether the *MK* lines continued to produce female-only offspring across generations, we also set up vials of *N101*^*MK*^ and *B302*^*MK*^ which were crossed to *B116*^−^ males each generation. We set up 5-10 replicate vials for each treatment and control and these were tracked and maintained separately. Each generation, lines that produced only females were maintained by adding 5 *B116*^−^ males, otherwise males were not added. Lines were allowed to mate for 3 days and were then transferred to new food to produce offspring for 4 days. Lines were scored for sex ratio every generation and maintained for 5 generations.

## Acknowledgements

The authors would like to thank Qiong Yang, Katie Robinson, Nancy Endersby-Harshman and Tim Wheeler for molecular assistance. We also thank Jake Brown, Jackson Young, Courtney Brown, Torsten Kristensen, Andres Andersen and Christian Danielsen for technical assistance.

## Supporting Information

**Figure S1. Relative densities of the (A) CI and (B) MK *Wolbachia* strains in *Drosophila pseudotakahashii* lines after long-term laboratory culture**. Data points show densities in individual adults while vertical lines and error bars show medians and 95% confidence intervals. Individuals testing negative for a *Wolbachia* strain were excluded.

**Figure S2. Relative densities of the (A, C) CI and (B, D) MK *Wolbachia* strains in *Drosophila pseudotakahashii* lines following backcrossing**. Females from the (A-B) *N101*^*MKS*^ line or (C-D) *B302*^*MKS*^ line were crossed to *B116*^−^ males for three generations. *Wolbachia* density was measured in the original lines and backcrossed lines that produced both male and female offspring (*MKS*) or female-only offspring (*MK*). Data points show densities in individual adults while vertical lines and error bars show medians and 95% confidence intervals. Individuals testing negative for a *Wolbachia* strain were excluded.

**Table S1**. Infection types present in samples screened from lines originally expressing CI and MK phenotypes at F11

**Table S2**. Private allele counts, observed heterozygosity (H_O_), expected heterozygosity (H_E_), and inbreeding coefficients (F_IS_) for the female-only and mixed-sex phenotypes of the N101 and B302 lines.

**Table S3**. Collection locations for *D. pseudotakahashii* populations used in this study.

## Notes

### Competing Interest Statement

The authors have declared no competing interest.

## References

1. Majerus M, Hurst G. Ladybirds as a model system for the study of male-killing symbionts. Entomophaga. 1997;42(1):13–20.

2. Hurst GDD, Majerus MEN. Why do maternally inherited microorganisms kill males? Heredity. 1993;71:81–95. doi: 10.1038/hdy.1993.110.

3. Jiggins FM, Randerson JP, Hurst GDD, Majerus MEN. How can sex ratio distorters reach extreme prevalences? Male-killing Wolbachia are not suppressed and have near-perfect vertical transmission efficiency in Acraea encedon. Evolution. 2002;56(11):2290–5.

4. Dyer KA, Jaenike J. Evolutionarily Stable Infection by a Male-Killing Endosymbiont in Drosophila innubila: Molecular Evidence From the Host and Parasite Genomes. Genetics. 2004;168(3):1443–55. doi: 10.1534/genetics.104.027854.

5. Dyer KA, Minhas MD, Jaenike J. Expression and modulation of embryonic male-killing in Drosophila innubila: opportunities for multilevel selection. Evolution. 2005;59(4):838–48. doi: 10.1111/j.0014-3820.2005.tb01757.x.

6. Richardson KM, Schiffer M, Griffin PC, Lee SF, Hoffmann AA. Tropical Drosophila pandora carry Wolbachia infections causing cytoplasmic incompatibility or male killing. Evolution. 2016;70(8):1791–802.

7. Montenegro H, Hatadani LM, Medeiros HF, Klaczko LB. Male killing in three species of the tripunctata radiation of Drosophila (Diptera: Drosophilidae). J Zool Syst and Evol Res. 2006;44(2):130–5.

8. Sheeley SL, McAllister BF. Mobile male-killer: similar Wolbachia strains kill males of divergent Drosophila hosts. Heredity. 2009;102(3):286–92. doi: 10.1038/hdy.2008.126.

9. Mateos M, Castrezana SJ, Nankivell BJ, Estes AM, Markow TA, Moran NA. Heritable endosymbionts of Drosophila. Genetics. 2006;174(1):363–76.

10. Nguyen DT, Morrow JL, Spooner-Hart RN, Riegler M. Independent cytoplasmic incompatibility induced by Cardinium and Wolbachia maintains endosymbiont coinfections in haplodiploid thrips populations. Evolution. 2017;71(4):995–1008.

11. Vala F, Breeuwer JA, Sabelis MW. Wolbachia–induced ‘hybrid breakdown’ in the two–spotted spider mite Tetranychus urticae Koch. Proc Biol Sci. 2000;267(1456):1931–7.

12. Jaenike J, Dyer KA. No resistance to male-killing Wolbachia after thousands of years of infection. J of Evol Biol. 2008;21(6):1570–7. doi: 10.1111/j.1420-9101.2008.01607.x.

13. Hornett EA, Duplouy AM, Davies N, Roderick GK, Wedell N, Hurst GD, et al. You can’t keep a good parasite down: Evolution of a male-killer suppressor uncovers cytoplasmic incompatibility. Evolution. 2008;62(5):1258–63.

14. Hornett EA, Engelstädter J, Hurst GDD. Hidden cytoplasmic incompatibility alters the dynamics of male-killer/host interactions. J of Evol Biol. 2010;23(3):479–87. doi: 10.1111/j.1420-9101.2009.01872.x.

15. Dyson EA, Hurst GD. Persistence of an extreme sex-ratio bias in a natural population. PNAS. 2004;101(17):6520–3.

16. Hornett EA, Charlat S, Duplouy AMR, Davies N, Roderick GK, Wedell N, et al. Evolution of Male-Killer Suppression in a Natural Population. PLoS Biol. 2006;4(9):e283.

17. Hayashi M, Nomura M, Kageyama D. Rapid comeback of males: evolution of male-killer suppression in a green lacewing population. Proc Biol Sci. 2018;285(1877):20180369.

18. Yoshida K, Sanada-Morimura S, Huang S-H, Tokuda M. Silence of the killers: discovery of male-killing suppression in a rearing strain of the small brown planthopper, Laodelphax striatellus. Proc Biol Sci. 2021;288(1943):20202125.

19. Reynolds LA, Hornett EA, Jiggins CD, Hurst GD. Suppression of Wolbachia-mediated male-killing in the butterfly Hypolimnas bolina involves a single genomic region. PeerJ. 2019;7:e7677.

20. Jaenike J. Spontaneous emergence of a new Wolbachia phenotype. Evolution. 2007;61(9):2244–52.

21. Richardson KM, Griffin PC, Lee SF, Ross PA, Endersby-Harshman NM, Schiffer M, et al. A Wolbachia infection from Drosophila that causes cytoplasmic incompatibility despite low prevalence and densities in males. Heredity. 2019;122(4):428–40.

22. Li F, Rane RV, Luria V, Xiong Z, Chen J, Li Z, et al. Phylogenomic analyses of the genus Drosophila reveals genomic signals of climate adaptation. Mol Ecol Resour. 2022;22:1559–81.

23. Baldo L, Hotopp JCD, Jolley KA, Bordenstein SR, Biber SA, Choudhury RR, et al. Multilocus sequence typing system for the endosymbiont Wolbachia pipientis. Appl Environ Microbiol. 2006;72(11):7098–110.

24. Turelli M, Hoffmann AA,. Cytoplasmic incompatibility in Drosophila simulans: dynamics and parameter estimates from natural populations. Genetics. 1995;140(4):1319–38.

25. Hurst GD, Jiggins FM, Robinson SJ. What causes inefficient transmission of male-killing Wolbachia in Drosophila? Heredity. 2001;87(2):220–6.

26. LePage DP, Metcalf JA, Bordenstein SR, On J, Perlmutter JI, Shropshire JD, et al. Prophage WO genes recapitulate and enhance Wolbachia-induced cytoplasmic incompatibility. Nature. 2017;543(7644):243–7.

27. Beckmann JF, Ronau JA, Hochstrasser M. A Wolbachia deubiquitylating enzyme induces cytoplasmic incompatibility. Nat Microbiol. 2017;2(5):1–7.

28. Shropshire JD, Hamant E, Conner WR, Cooper BS. cifB-transcript levels largely explain cytoplasmic incompatibility variation across divergent Wolbachia. PNAS Nexus. 2022;1(3):pgac099. doi: 10.1093/pnasnexus/pgac099.

29. Martinez J, Klasson L, Welch JJ, Jiggins FM. Life and Death of Selfish Genes: Comparative Genomics Reveals the Dynamic Evolution of Cytoplasmic Incompatibility. Mol Biol and Evol. 2020;38(1):2–15. doi: 10.1093/molbev/msaa209.

30. Poinsot D, Bourtzis K, Markakis G, Savakis C, Merçot H. Wolbachia Transfer from Drosophila melanogaster into D. simulans: Host Effect and Cytoplasmic Incompatibility Relationships. Genetics. 1998;150(1):227–37. doi: 10.1093/genetics/150.1.227.

31. Ellegaard KM, Klasson L, Näslund K, Bourtzis K, Andersson SGE. Comparative Genomics of Wolbachia and the Bacterial Species Concept. PLoS Genet. 2013;9(4):e1003381. doi: 10.1371/journal.pgen.1003381.

32. Turelli M, Cooper BS, Richardson KM, Ginsberg PS, Peckenpaugh B, Antelope CX, et al. Rapid global spread of wRi-like Wolbachia across multiple Drosophila. Curr Biol. 2018;28(6):963-71. e8.

33. Cooper BS, Vanderpool D, Conner WR, Matute DR, Turelli M. Wolbachia acquisition by Drosophila yakuba-clade hosts and transfer of incompatibility loci between distantly related Wolbachia. Genetics. 2019;212(4):1399–419.

34. Gerth M, Bleidorn C. Comparative genomics provides a timeframe for Wolbachia evolution and exposes a recent biotin synthesis operon transfer. Nat Microbiol. 2016;2(3):1–7.

35. Klasson L, Walker T, Sebaihia M, Sanders MJ, Quail MA, Lord A, et al. Genome evolution of Wolbachia strain wPip from the Culex pipiens group. Mol Biol Evol. 2008;25(9):1877–87.

36. Meany MK, Conner WR, Richter SV, Bailey JA, Turelli M, Cooper BS. Loss of cytoplasmic incompatibility and minimal fecundity effects explain relatively low Wolbachia frequencies in Drosophila mauritiana. Evolution. 2019;73(6):1278–95.

37. Bock I, Parsons P. Australian endemic Drosophila IV. Queensland rain forest species collected at fruit baits, with descriptions of two species. Aust J Zool. 1978;26(1):91–103.

38. Hague MT, Shropshire JD, Caldwell CN, Statz JP, Stanek KA, Conner WR, et al. Temperature effects on cellular host-microbe interactions explain continent-wide endosymbiont prevalence. Curr Biol. 2022;32(4):878-88. e8.

39. Ross PA, Wiwatanaratanabutr I, Axford JK, White VL, Endersby-Harshman NM, Hoffmann AA. Wolbachia infections in Aedes aegypti differ markedly in their response to cyclical heat stress. PLoS Pathog. 2017;13(1):e1006006.

40. Rousset F, Solignac M. Evolution of single and double Wolbachia symbioses during speciation in the Drosophila simulans complex. PNAS. 1995;92(14):6389–93.

41. Hornett EA, Kageyama D, Hurst GD. Sex determination systems as the interface between male-killing bacteria and their hosts. Proc Biol Sci. 2022;289(1972):20212781.

42. Hurst GDD, Jiggins FM. Male-killing bacteria in insects: Mechanisms, incidence, and implications. Emerg Infect Dis. 2000;6(4):329–36. PubMed PMID: WOS:000088660900002.

43. Jaenike J, Dyer K. No resistance to male-killing Wolbachia after thousands of years of infection. J Evol Biol. 2008;21(6):1570–7.

44. O’Neill SL, Hoffmann A, Werren J. Influential passengers: inherited microorganisms and arthropod reproduction: Oxford University Press; 1997.

45. Bock IR. Drosophilidae of Australia. I. Drosophila (Insecta: Diptera). Aust J Zool Supp. 1976;24(40):1–105.

46. Matsuda M, Ng C-S, Doi M, Koppa A, Toba YN. Evolution in the Drosophila ananassae species subgroup. Fly. 2009;3(2):1–13.

47. Folmer O, Black M, Hoeh W, Lutz R, Vrijenhoek R. DNA primers for PCR amplification of mitochondrial cytochrome c oxidase subunit I from diverse metazoan invertebrates. Mol Mar Biol and Biotechnol. 1994;3(5):294–9.

48. Tamura K, Stecher G, Peterson D, Filipski A, Kumar S. MEGA6: molecular evolutionary genetics analysis version 6.0. Mol Biol Evol. 2013;30(12):2725–9.

49. Hoffmann AA, Turelli M, Simmons GM. Unidirectional incompatibility between populations of Drosophila simulans. Evolution. 1986;40:692–701.

50. Kriesner P, Hoffmann AA, Lee SF, Turelli M, Weeks AR. Rapid Sequential Spread of Two Wolbachia Variants in Drosophila simulans. PLOS Path. 2013;9(9):e1003607.

51. Lee SF, White VL, Weeks AR, Hoffmann AA, Endersby NM. High-throughput PCR assays to monitor Wolbachia infection in the dengue mosquito (Aedes aegypti) and Drosophila simulans. Appl Environ Microbiol. 2012;78(13):4740–3.

52. Bleidorn C, Gerth M. A critical re-evaluation of multilocus sequence typing (MLST) efforts in Wolbachia. FEMS Microbiol Ecol. 2018;94(1):fix163.

53. Walker BJ, Abeel T, Shea T, Priest M, Abouelliel A, Sakthikumar S, et al. Pilon: an integrated tool for comprehensive microbial variant detection and genome assembly improvement. PloS One. 2014;9(11):e112963.

54. Simão FA, Waterhouse RM, Ioannidis P, Kriventseva EV, Zdobnov EM. BUSCO: assessing genome assembly and annotation completeness with single-copy orthologs. Bioinformatics. 2015;31(19):3210–2.

55. Wu M, Sun LV, Vamathevan J, Riegler M, Deboy R, Brownlie JC, et al. Phylogenomics of the reproductive parasite Wolbachia pipientis wMel: a streamlined genome overrun by mobile genetic elements. PLoS Biol. 2004;2(3):e69.

56. Li H, Durbin R. Fast and accurate short read alignment with Burrows–Wheeler transform. Bioinformatics. 2009;25(14):1754–60.

57. Shropshire JD, On J, Layton EM, Zhou H, Bordenstein SR. One prophage WO gene rescues cytoplasmic incompatibility in Drosophila melanogaster. PNAS. 2018;115(19):4987–91.

58. Beckmann JF, Van Vaerenberghe K, Akwa DE, Cooper BS. A single mutation weakens symbiont-induced reproductive manipulation through reductions in deubiquitylation efficiency. PNAS. 2021;118(39):e2113271118.

59. Perlmutter JI, Bordenstein SR, Unckless RL, LePage DP, Metcalf JA, Hill T, et al. The phage gene wmk is a candidate for male killing by a bacterial endosymbiont. PLoS Path. 2019;15(9):e1007936.

60. Choi JY, Bubnell JE, Aquadro CF. Population genomics of infectious and integrated Wolbachia pipientis genomes in Drosophila ananassae. Genome Biol Evol. 2015;7(8):2362–82.

61. Sutton ER, Harris SR, Parkhill J, Sinkins SP. Comparative genome analysis of Wolbachia strain wAu. BMC Genomics. 2014;15(1):1–15.

62. Klasson L, Westberg J, Sapountzis P, Näslund K, Lutnaes Y, Darby AC, et al. The mosaic genome structure of the Wolbachia wRi strain infecting Drosophila simulans. PNAS. 2009;106(14):5725–30.

63. Seemann T. Prokka: rapid prokaryotic genome annotation. Bioinformatics. 2014;30(14):2068–9.

64. Rašic G, Filipovic I, Weeks AR, Hoffmann AA. Genome-wide SNPs lead to strong signals of geographic structure and relatedness patterns in the major arbovirus vector, Aedes aegypti. BMC Genomics. 2014;15(1):1–12.

65. Catchen J, Hohenlohe PA, Bassham S, Amores A, Cresko WA. Stacks: an analysis tool set for population genomics. Mol Ecol. 2013;22(11):3124–40.

66. Langmead B, Salzberg SL. Fast gapped-read alignment with Bowtie 2. Nat Methods. 2012;9(4):357–9.

67. Kim BY, Wang JR, Miller DE, Barmina O, Delaney E, Thompson A, et al. Highly contiguous assemblies of 101 drosophilid genomes. eLife. 2021;10:e66405. doi: 10.7554/eLife.66405.

68. Schmidt TL, Jasper ME, Weeks AR, Hoffmann AA. Unbiased population heterozygosity estimates from genome-wide sequence data. Methods Ecol Evol. 2021;12(10):1888–98.

69. Linck E, Battey C. Minor allele frequency thresholds strongly affect population structure inference with genomic data sets. Mol Ecol Resour. 2019;19(3):639–47.

